# Discovering Secondary Protein Structures via Local Euler Curvature

**DOI:** 10.1101/2023.11.27.568841

**Authors:** Rodrigo A. Moreira, Roisin Braddell, Fernando A. N. Santos, Tamàs Fülöp, Mathieu Desroches, Iban Ubarretxena-Belandia, Serafim Rodrigues

## Abstract

Protein structure analysis and classification, which is fundamental for predicting protein function, still poses formidable challenges in the fields of molecular biology, mathematics, physics and computer science. In the present work we exploit recent advances in computational topology to define a new intrinsic unsupervised topological fingerprint for proteins. These fingerprints, computed via Local Euler Curvature (LECs), identify secondary protein structures, such as Helices and Sheets, by capturing their distinctive topological signatures. Using an extensive protein residue database, the proposed computational framework not only distinguishes between structural classes via unsupervised clustering but also achieves remarkable accuracy in classifying proteins structures through supervised machine learning classifier. We also show that the internal structure of LEC space embeds the information about the secondary structure of proteins. Beyond its immediate implications for the advancement of critical application areas such as drug design and biotechnology, our approach opens a fascinating avenue towards characterizing the multiscale structures of diverse biopolymers based solely on their geometric and topological attributes.

---

Proteins are complex biomolecules that assemble cellular organisms into life by orchestrating cellular structure, energy and information processing. These vital functions include for example, structural support, enzymatic reactions, signal transduction and beyond, which together lead to emergent cellular processes. The molecular mechanisms of protein functions are closely related to their structures. Thus, predicting their three-dimensional (3D) structure-functional relationship is the ultimate goal of biophysics. Predictions will lead to significant advances in drug design, medicine, biotechnology and the understanding of cellular life. Traditional methods for determining protein structures involve experimental approaches like X-ray crystallography, NMR spectroscopy and circular dichroism.^1^. Limits of experimental protocols has led to the development of algorithms to classify proteins. Commonly used protein secondary structure assignment^2^ include the DSSP^3^ and STRIDE^4^ algorithms. These procedures are based on a hydrogen-bond^5^ (hbond) distance and/or on an energetic cutoff between the C=O and N-H groups present in a protein’s backbone or previous knowledge about the protein structure. Both classifications schemes use molecular models and experiments as tools to standardize the assignment of secondary protein structures into roughly seven groups: The turns (T) and coils (C); The three helix types, the 3_10_-Helix (G), α-Helix (H) and *τ*-Helix (I), in which the residues are repeated after three, four, and five units, respectively, and also two kinds of extended structures, namely, a,*B*-Strand (E) and a,*B*-Bridge(B). Some of these structures are represented in Fig. 1a, where for convenience we depict some distances related to each secondary structure.

**Fig. 1.**
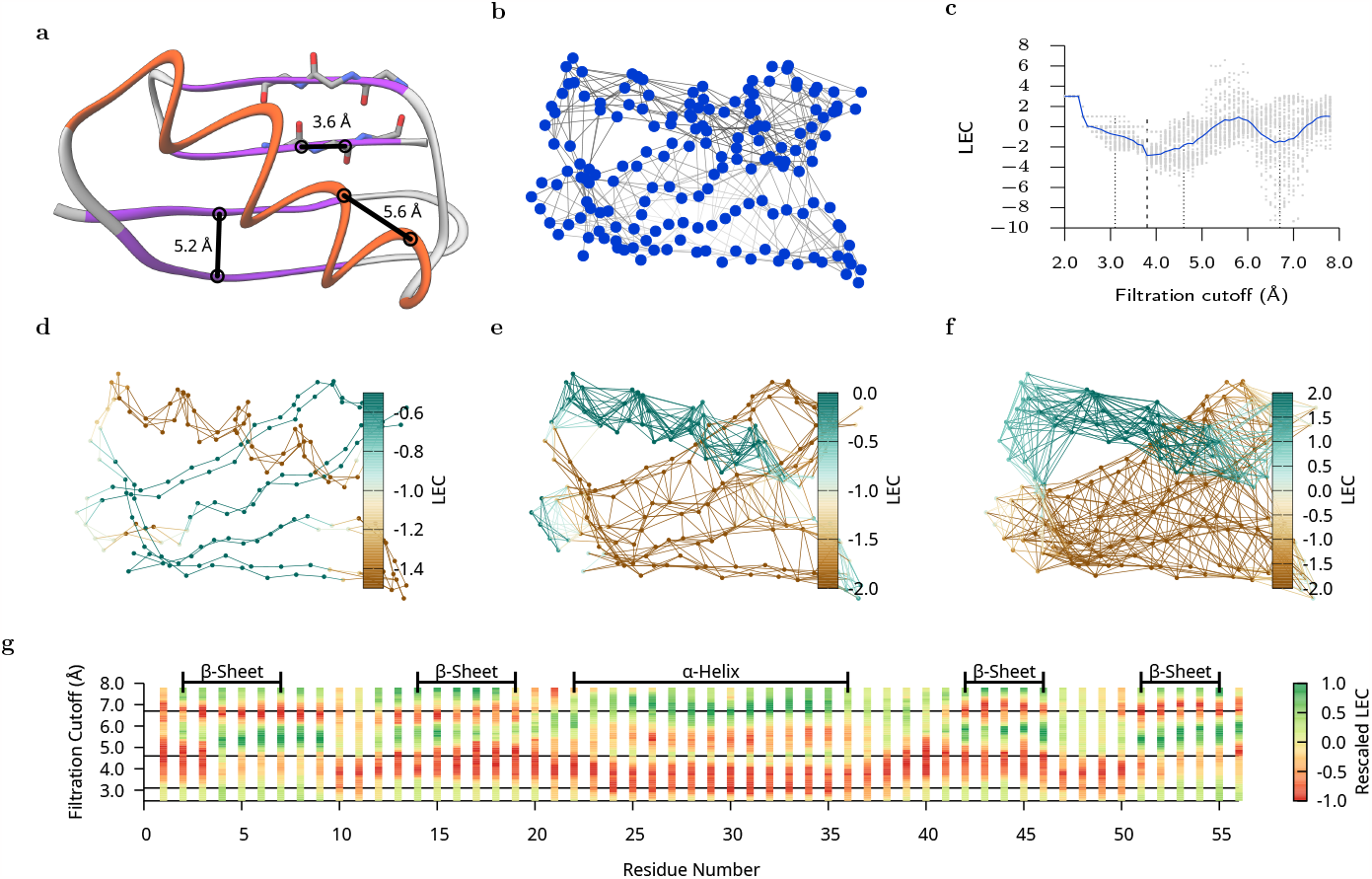
Representations of protein PDB 2GB1. **(a)** We highlight the relevant distances. Typical approximate distances 3.6 Å for (53C-54C) backbone length and (5CA-16CA) 5.2 Å and (31CB-34CB) 5.6 Å for,*B*-Strand interchain separation and *α*-Helix, respectively. The colors purple/orange/grey represent the standard DSSP assignments of,*B*-Strand, *α*-Helix and Turns/Coils, respectively. **(b)** Graph representation showing the nodes in blue and the edges in grey.**(c)** LEC representation, where grey dots are the computed Local Euler Curvature as a function of the cutoff distance of all residues from 2GB1 and the blue line is its average. **(d)-(f)** Representation at different important filtration cutoffs 3.1, 4.6 and 6.7 Å(also indicated in Fig. 3b) of the 0-simplices and 1-simplices, where the colors represent LEC (Eq. 4) of each residue accordingly to respective colorbars. **(g)** LEC profiles for all residue of 2GB1, where we also indicate by solid horizontal lines the 3 cutoffs 3.1, 4.6, 6.7Å. The LEC profiles were re-scaled between [-1:1] to improve the contrast.

More recently, machine learning approaches have significantly advanced protein structural analysis research^6–11^. Among these, the most impressive is the Google DeepMind’s AlphaFold AI program^12^, which has won the Casp and Lasker basic medical research awards for predicting (with circa 95% accuracy) protein’s 3D structures from their amino-acid sequences. Despite these advances, machine learning models are essentially black-boxes, thus they fail to explain the biophysical mechanisms associated with protein’s multi-scale structures, intricate folded patterns,^13^, interactions^14^ or interfaces.^15–18^. Consequently, a parallel line of research has been exploring more mathematical approaches based on geometry and topology. This has been enabled by recent advances in computational topology that provide efficient algorithms to synthesise highly complex nonlinear high-dimensional data as geometrical objects (‘shapes’), which satisfy properties (called *Topological invariants*) remaining unchanged under continuous deformations (e.g. stretching, twisting, or bending)^19–24^. With this approach, data is represented via *Simplicial Complexes (SC)*^25^, which generalize the notion of triangulation of a surface and are constructed by gluing together *simplices* (i.e. points, line segments, triangles and their higher dimensional counterparts). The gluing satisfy certain nested conditions (i.e. higher simplexes must contain lower simplexes). The gluing or reconstruction of the data typically follows a combinatorial process (e.g. *Simplicial Filtration*, sometimes referred as *Filtration cutoff* ) endowed with a distance function, which then constructs a family (indexed by distance) of simplicial complexes (i.e. growing sequences of meshes). Every simplicial complex in this sequence is then studied via topological invariants (e.g. *peristent Barcodes, Euler-Poincaré characteristics* etc). Applications of computational topology to the study of proteins include, protein structures analysis^26–28^, molecular interactions in the context of molecular simulations^29–31^, and also hybrid approaches combining machine learning methods with computational topology^32–34^. Noteworthy, some hybrid approaches have achieved 85% accuracy in protein classification^32^ and notably the MathDL hybrid algorithm has won the D3R Grand Challenge 4^35^. Despite these advances, a direct classification of protein structures from their topology is still an open problem. Indeed, to date, most computational topological approaches employ *Barcode* as the main topological invariant since it is *complete*^36^ and is endowed with *stability conditions*^37^. However, the Space of all barcodes is not a vector space and as a consequence is not directly amenable to analysis (e.g. statistical analysis), although there are ongoing theoretical developments in this direction (e.g. *Persistent landscapes*^38^, *barcode stratification via Coxeter complexes*^39^). In short, barcodes contain excessive information, does not lend itself to analysis and interpretation of results, which leads to difficulties in protein classification.

To advance, we take the advantage of a key mathematical property of the *Euler characteristics* (EC), which is that it can be decomposed (or partitioned), leading to the concept of *Local Euler Curvature*, or in this work sometimes called also Local Euler Characteristics, (LEC)^40–42^. LEC lends itself to protein analysis since local properties of proteins are crucial. Overall, this leads to a more efficient compartmentalised analysis and interpretation, while still connecting to the global topological properties of proteins. Specifically, we will show that the basic secondary structure of proteins are naturally described by LEC. To validate this, we analysed a representative database containing millions of protein residues, and subsequently modelled and scored using LEC representation. Importantly, we show for the first time, that the structures of proteins can be precisely classified based only on their topology, using a simple methodology based on LEC representation. We organise this sequel as follows; we first reconstruct a sequence of simplicial complexes (i.e. via filtration) and compute their corresponding LEC for a specific protein, which provides an overview of the computational pipeline. We then apply the LEC in the context of an ensemble of proteins to maximise the mapping of secondary structures and compare with previous classification schemes. These fundamental findings is then used in a unsupervised learning framework to show how secondary structures can be mapped and classified via LEC. Moreover, we employ a *Random Forest Classifier* model to predict the proteins secondary structures relying solely on their reduced backbone and respective topology. Finally, we conclude this study with a discussion section, which also outlines future perspectives.

## Results

As a brief summary of what results are to be expected: We will first show in Fig. 1 that LEC of protein residues naturally describes the traditional secondary structure assignment from standard methodologies. Then, these qualitative results will be generalized to an ensemble of protein structures (see Fig. 2), and the individual LEC signatures of 7 secondary structure classes will be characterized by their LEC profiles. Once the robust statistics of our aggregation of classes into different LEC filtration have been established, we will reinterpret the filtration process as a topological feature space, which will be used for learning the protein structures in an unsupervised manner. Additionally, we will train a *Random Forest Classifier* model and subsequently its final scores in comparison to reference methodologies will be analysed in Table 1.

**Table 1.** Final scores of our data sets. The number of residues used was 17,875 at final assessment test set and 2,215,533 for our CATH data set. f1-score is defined by *f* 1 = 2(precision *\** recall)/(precision + recall).

| Protein |  | f1-score |  |  |  | Accuracy |  |
| --- | --- | --- | --- | --- | --- | --- | --- |
| Prediction | Reference | β-Strand | α-Helix | Turn | Coil | Score | Top$K=2$ |
| Test data set |  |  |  |  |  |  |  |
| rbLEC | DSSP | 0.86 | 0.93 | 0.52 | 0.79 | 0.79 | 0.90 |
| rbLEC | STRIDE | 0.79 | 0.91 | 0.58 | 0.69 | 0.74 | 0.87 |
| STRIDE | DSSP | 0.86 | 0.95 | 0.44 | 0.69 | 0.76 | - |
| CATH data set |  |  |  |  |  |  |  |
| rbLEC | DSSP | 0.87 | 0.94 | 0.60 | 0.80 | 0.81 | 0.91 |
| rbLEC | STRIDE | 0.70 | 0.91 | 0.59 | 0.62 | 0.71 | 0.84 |
| STRIDE | DSSP | 0.75 | 0.93 | 0.46 | 0.62 | 0.72 | - |

**Fig. 2.**
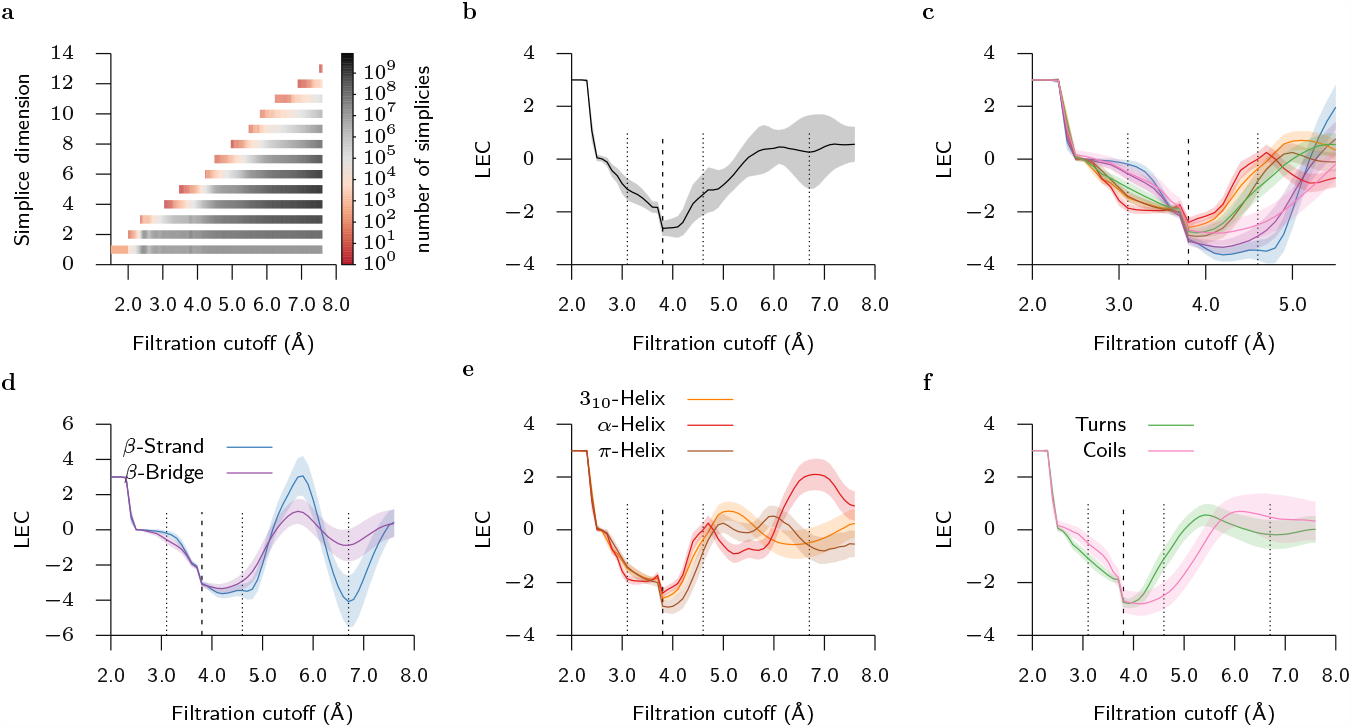
Topology of the consensus residues from CATH database. **(a)** Average number of simplices (#simplices) of all molecules vs filtration cutoff. Higher than 13-simplices were not found in this range. **(b)-(f)** LEC computed using (Eq. (4)) from all consensus residues. The average and standard deviations are represented by the solid line and filled curves, respectively. In **b** we show the average over all consensus residues, while the indicated colours in **(c)** to **(f)** represent different consensus secondary structures. Vertical lines show relevant distances, where the bold line represents typical 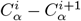 distance, and dotted lines are the distances highlighted in Fig. 3b.

### Local Euler Curvature of Proteins

We start our exploratory work by studying a specific example, namely the protein 2GB1, which will guide the reminder of our study on an ensemble of proteins. We first illustrate, in Fig. 1a, the conventional process of assigning secondary structures to the protein 2GB1 using the DSSP algorithm. We then proceed via a computational combinatorial topology approach where a discrete topological space is associated to the 2GB1 protein structure via a *Simplicial Filtration* approach (see Methods). Such a construction leads to the simplicial complex (SC) depicted in Fig. 1b, for a fixed filtration cutoff parameter. However, Fig. 1b can be pruned to accommodate a specific maximum or minimum edge distance, hence yielding distinct SCs (i.e. filtration). For instance, Figs. 1d-1f showcase the respective SC for cutoff distances of 3.1 Å, 4.6 Å, and 6.7 Å, respectively, which also illustrates the combinatorial explosion of edges that emerge as a function of increase cutoff. Moreover, for every generated SC, we compute its corresponding *Euler Characteristics and Local Euler Curvature* (see Methods). The LEC can be computed per atom of the protein, however this leads to a fine-grained computation, which is unnecessary for residue-based protein structural classification. Thus, we restrict to the backbone network, comprising PDB atoms CA, C, N of each protein residue. Hereafter, the network used to compute the simplical simplex is formed by its restricted backbone and the LEC of a residue is given by PDB atoms CA, C and N (see Methods). Such computations is shown in Fig. 1c, which depicts that along the filtration process (within the cutoff range 1.5 to 7.5 Å), for every generated SC, the corresponding LEC associated to every residue of 2GB is computed (grey dots), as well as, their average (blue curve). Strikingly, we observe that our SC representation and associated LEC is highly interpretable by relevant chemical distances. For example, two consecutive *C*^*α*^ of protein chains have a typical distance of approximately 4.0 Å, which corresponds to a minimum of LEC. Further examining the plots of LEC values for individual protein residues at various cutoff distances, as depicted in Figs. 1d-1f, we can easily visually identify the standard *α*-Helix and,*B*-Strand/,*B*-Bridge secondary structure of the protein. Noteworthy, this identification emerges due to the recognizable patterns of Local Euler Curvature.

However, proteins exhibit a whole spectrum of diverse shapes and sizes and as a consequence, this requires a more comprehensive analysis, which can only be captured in the study of en ensemble of proteins. This is critical if we are to establish a robust correlation between topological invariants and protein secondary structures.

### Ensemble Local Euler Curvature of Proteins

We considered a representative from the CATH database^43^, which maintains a collection of nonredundant protein structures; see Methods. In all proteins we first selected the three backbone atoms CA, C, and N as the nodes of our graph. Then following our approach from the preceding section, we computed the simplices and their corresponding local curvatures (specifically Eq. (1)), which were then aggregated to yield the LEC (Eq. (4)) for each individual residue. Our data aggregation encompassed a total of 2,215,533 residues.

Proceeding with the *Simplicial Filtration* approach by varying the cutoff distance from 1.5 to 7.5 Å we observe a combinatorial explosion in the emergence of simplices as depicted in Fig. 2a. Indeed, note the exponential growth, with an average of approximately 10^9^ simplices in dimensions 4 and 5 at a 7.5 Å cutoff. Within this cutoff range, the highest dimensional simplices were of dimension 14 and the most occurring ones were of dimensions 4, 5, and 6.

We then followed by computing the LEC of the generated simplices. Fig. 2b depicts the average LEC of the residues, along with their respective standard deviations, across varying cutoff distances. Remarkably, a close correspondence between the averages of LEC in Fig. 1c and those in Fig. 2b is discernible. Despite the former originating from a single structure comprising 56 residues and the latter stemming from a large ensemble of residues, this alignment underscores the robustness of our methodology.

Given the distinct secondary structures inherent in proteins, we decided to focus on the seven categories elaborated upon in the *Introduction*, namely,*B*-Bridge,*B*-Strand, *α*-Helix, 3_10_-Helix, *τ*-Helix, Coils, and Turns. To delve deeper, we employed standard software to categorize residues into these seven classes, computing the individual averages for each. To this end, we exclusively considered residues with matching DSSP and STRIDE assignment, denoted as the *consensus* classification.

The results presented in Fig. 2c-2f, show the LEC averages and standard deviations after the consensus classification of CATH residues. Fig. 2c displays the average LEC across the seven secondary structure categories within the [1.5, 5.5] Å cutoff range. One key finding is a distinctive gap between extended structures (,*B*-Bridge,,*B*-Strand, and Coils) and highly turned structures (*α*), including *α*-Helix, 3_10_-Helix, *τ*-Helix, and Turns, emerged just beyond the 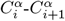 distance at around 4.6 Å. The differentiation between all classes was further evident in the vicinity of 3.1Å, notably highlighting the distinctiveness of *α*-Helix and,*B*-Strand structures, as previously seen in Figs. 1d-1f. Extending the analysis up to a 7.5 Å cutoff (Figs. 2d-2f), yields a finer differentiation across all seven categories.

In particular, the,*B* structures display a LEC *plateau* after 4.0 Å in Fig. 1d. The similarity between,*B*-Strand and,*B*-Bridge persist until 5.0 Å, after which,*B*-Bridge exhibit smaller oscillations than the characteristic,*B*-Strand oscillations, as indicated by the reduced standard deviation, including a gap around 6.7 Å.

In contrast, the *α* structures in Fig. 2e showcase various patterns. While the three helix types share similar LEC profiles, they do not feature the *plateau* characteristic of,*B* structures. Divergence among *α* structures initiate at 5.0 Å, where 3_10_-Helix and *τ*-Helix demonstrate predominantly positive LEC values between 5.0 and 6.0 Å, while *α*-Helix exhibit negative values. Subsequently, beyond 6.0 Å, *α*-Helix acquire positive values, while the other two exhibit negative values. Notably, the relative small standard deviation of LEC demonstrates remarkable homogeneity across a wide range of cutoff distances.

The behaviour of the remaining classes, Turns and Coils, is illustrated in Fig. 2f. Interestingly, they exhibit intermediate characteristics, with Turns resembling 3_10_-Helix trajectories, while Coils mostly resemble,*B*-Bridge structures. However, both categories display minimal oscillations beyond 6.0 Å, distinguishing them from what is shown on Figs. 2d and 2e.

The correlation between typical distances, such as *α*-Helix, pitch and,*B*-Strand interchain distances (Fig. 1), with the trends described in the previous sections, highlights the utility of our methodology in capturing plateaus and oscillations within distinct ranges, enhancing its versatility.

Up to this point, our results suggest that we can effectively identify *α* and,*B* structures in the scenario depicted in Fig. 1. Furthermore, we can provide detailed and nuanced descriptions for all seven structural categories as illustrated in Fig. 2. Henceforth, we shall show that the observed LEC features are sufficient to classify protein secondary structures. To this end, we employ an unsupervised clustering algorithm to reveal a clear classification of protein structures resembling those depicted in Fig. 2. Moreover, the accuracy of our methodology is demonstrated via Random Forrest Classifier, which validates the proposed approach.

### Unsupervised Clustering of LEC profiles

Following an *Unsupervised Clustering* approach (see Methods), we uncover intriguing patterns in the clusters formed by LEC profile. Since these profiles depend on the spatial shape of the protein and should exhibit stability under minor structural deformations, high-density clusters of curves indicate large subsets of proteins with very similar spatial structure. To analyze the inherent relationships between these profiles, we assigned to each profile a vector in 51-dimensional space, with each vector component corresponding to the LEC calculated for the protein at cutoffs from 2.5 Å to 7.5 Å in steps of 0.1 Å.

As anticipated, strong correlations emerge among these vectors. Principal component analysis (PCA) showed that approximately 80% of the variance concentrated within the first five Principal Components (PCs); see Fig. 3d. Additionally, we employed a Gaussian Mixture Model (GMM) to identify distinctive clusters.

**Fig. 3.**
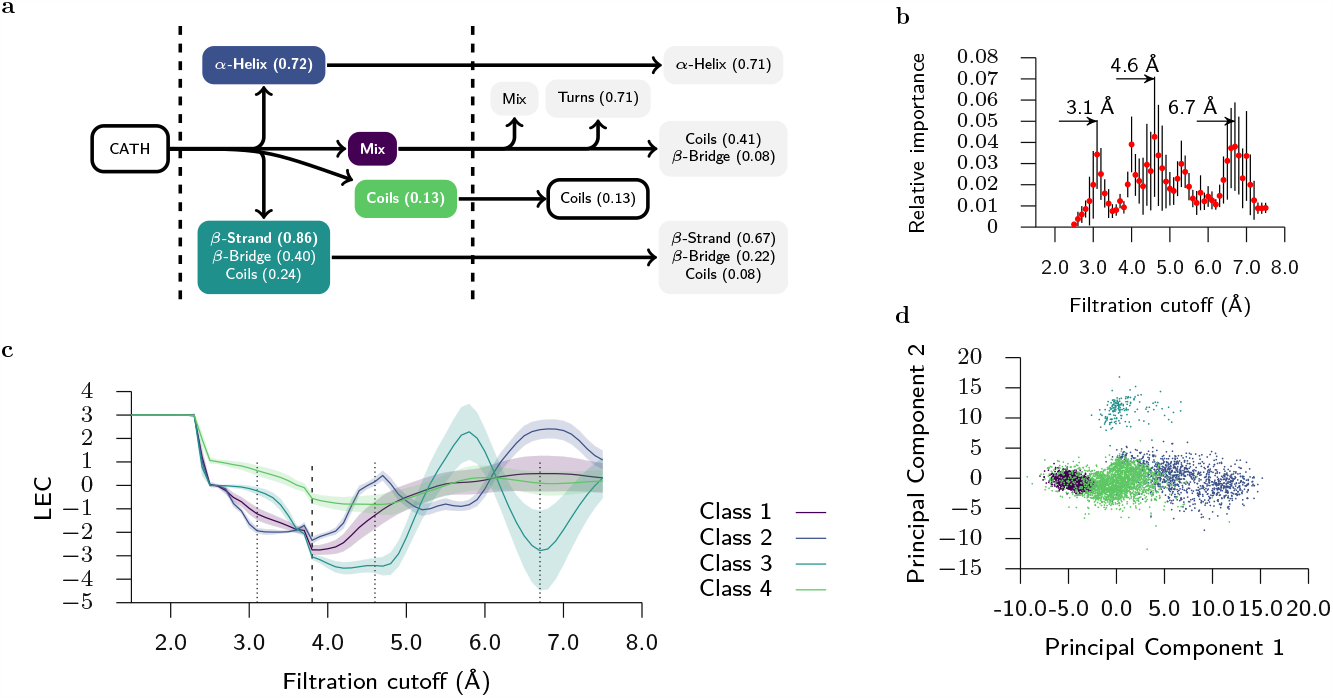
Machine learning LEC profiles. **(a)** Mostly hierarchy after 4 (left) and 6 (right) classes unsupervised clustering. **(b)** Importance for each filtration cutoff (features) returned by the random forest classifier. **(c)** Average (solid line) and standard deviation (filled area) of LEC profiles after unsupervised clustering in 4 classes. The vertical lines are the as in (Fig. 2). **(d)** Principal Components of **(c)**.

The GMM demonstrates a hierarchical structuring with six basic classes (see Fig. SI 11a for a density plot and Figs. SI 11c-SI 11f for a visualization of the clusters). These combine together reliably to form 3 or 4 clusters (see Fig. 3a for a schema of the relationships between 4 and 6 clusters detected by GMMs), which forms clearly distinguishable looking LEC profiles in Fig. 3c.

We tested the robustness of these clusterings via various measures outlined in Methods and Fig. SI 9. The broad characteristics of these clusters remain stable whether fitted on the full CATH dataset or the consensus subset. Noteworthy, Fig. SI 5 compared to Fig. SI 6 and Fig. SI 7, and also produce stable predictions when the GMM is fitted to a small portion of the dataset as described in Methods and Fig. SI 9.

Numerous combinations of those “basic classes” can arise as combinations if we select a different number of clusters, which leads to several different GMM fits in these cases. However, the classes demonstrate some expected results, such as dense clustering of *α*-Helix points (see Fig. SI 11a) and also include some surprising classes such as a cluster of about 13% of Coils; shown in Figs. SI 5 and SI 11c.

### Accuracy of LEC representation

The final findings we report here, focuses on evaluating how well our *Random Forest Classifier*, (denoted *rbLEC* ) (see Methods), performs when compared to traditional methods for assigning protein secondary structures. Table 1 displays the results in terms of f1-score and accuracy when compared to DSSP and STRIDE classifications.

First, it is important to gauge the accuracy of DSSP and STRIDE themselves, considering that they assign different classifications. We find that they have an accuracy of 76% for the CATH dataset (including non-consensus residues) and 72% for the test set. In contrast, our rbLEC model was trained exclusively on the consensus classification, omitting residues when DSSP and STRIDE do not agree. Thus, a comparable (possibly similar) scores between our rbLEC classifier and the DSSP/STRIDE is to be expected. Indeed, Table 1 shows that our rbLEC achieves a test set accuracy of 79% and 74% concerning DSSP and STRIDE, respectively, while CATH dataset yields 81% and 71%. Moreover, our rbLEC model provides class probabilities for a given feature. This allows us to compute the Top_*K*_ =2 accuracy, which is considered successful if the first or second most probable class matches the reference classification labels. We obtained scores of 90% and 87% for the test and CATH datasets, respectively. This remarkable result highlights the potential effectiveness of rbLEC and our training approach in providing a robust residue classifier for seven classes, based solely on the LEC of a given protein chain’s three backbone atoms. Notably, our methodology requires no *ad-hoc* determinations of hydrogen bond networks or protein geometric backbone geometries.

Additionally, Table 1 provides the f1-score for the four primary classes (encompassing over 95% of available residues):,*B*-Strand, *α*-Helix, Turns, and Coils. This score provides more information about the precision and recall balance of predicted classes. Interestingly, our rbLEC classifier yields f1-scores comparable to those of DSSP/STRIDE, corroborating our findings in the preceding sections.

## Discussion

Our model’s accuracy, as well as its performance in identifying the top two candidates, as shown in Table 1, is truly encouraging. To gain further insights and understand potential limitations, we turn to the confusion matrix presented in Table SI 2. This matrix, which predominantly features diagonal elements, suggests that our model excels in correctly classifying structures. However, it is worth noting that,*B*-Bridge and *τ*-Helix are the primary sources of relative errors.

Next, we dive into the interpretability of our methodology. We focus on,*B*-Bridge structures, which are often misclassified as,*B*-Strand, or Coils. From a structural and geometrical standpoint, these misclassifications are quite understandable. A closer examination of Fig. 2d and 2f reveals significant similarities in their LEC profiles, with noticeable differences only emerging after the 5.0 Å mark. In reality, finding perfectly flawless,*B*-Bridge conformations, as seen in isolated structures, is a rare occurrence in densely packed proteins.

What truly sets our methodology apart is its ability to automatically capture and classify structures, relying on the local environments of each residue. This is in contrast to conventional classification methods that rely on geometric rules, such as hydrogen bonding or residue counts. This innovative approach highlights the unique aspects of our method compared to traditional classification methods.

From our analysis, it is evident that Coils and Turns structures exhibit transitional features in their LEC profiles between helices and sheets, see Fig. SI 11b. Moreover, helices and sheets form distinct clusters when Coils and Turns are not considered. For other numbers of clusters, however, some clustering metrics markedly decrease and the Gaussians assigned by the GMM indicate higher variability. This may be due to the fact that a high number of new distinct cluster structures emerge in the data (see Fig. SI 12) that do not combine in a reliable way. Nevertheless, these clearly defined clusters open new interesting avenues of investigation.

We anticipate that a detailed examination of these categories will provide novel insights about protein secondary structures, shedding light onto cases where the classification systems do not align and potentially revealing interesting subtypes.

Assessing *τ*-Helix structures poses its own set of challenges due to limited consensus classifications in our test set (only 5 samples, see Table SI 1). Adding to the complexity are significant differences between the DSSP and STRIDE methodologies. From a geometric perspective, *τ*-Helix structures represent helices with an expanded radius, often nested within *α*-Helix, motifs. This arrangement introduces interruptions in the continuity of helices, potentially carrying significant functional implications for the protein. To gain deeper insights, future studies should explore these intertwined *α*-Helix structures using a more extensive dataset to establish a more robust consensus on *τ*-Helix classifications. Interestingly, our classifier demonstrates more consistent classification compared to DSSP than STRIDE, suggesting a potentially more uniform approach from DSSP authors theirs parameter choices.

In summary, the Euler characteristics (a global topological invariance) and its associated Local Euler Curvature (a local geometric measure) provides a powerful mathematical and computational tool to dissect in a consistent way the local and global properties associated to the topological representation of proteins. Specifically, we find t hat t he L ocal E uler C urvature p rovides a novel tool to classify protein’s secondary structures, which matches the state-of-the-art, however, without making strong geometrical assumptions. We envisage that our methodology will be useful to understand not only protein structures but also protein interactions in the context of disease models. For instance, this could be applied to Alzheimer’s Disease where it has been found that amyloid-beta peptides have anti-microbial role and may interact with glyco-proteins of various pathogens^44–48^. Moreover, we envisage that extensions of our proposed methodology will in the long-term contribute to the ambitious goal of predicting the 3D structure-functional relationship of proteins.

## Methods

### Data Collection and Preprocessing

Our dataset comprises 3D protein structures. The training set includes structures sourced from the latest version of the CATH database,^43, 49^ which categorizes protein structures from the Protein Data Bank into non-redundant domains. This compilation of nonredundant protein domains contains 14,938 files, representing 2,200,595 residues.

For the testing dataset, we acquired 22 proteins with a total of 17,875 residues from RCSB PDB.^50^ To ensure fairness, we selected a diverse range of complete proteins, encompassing both small (161 residues) and large (3797 residues) structures. This set encompasses prevalent tertiary structure groups such as *α* +,*B, α*/,*B*, all-*α*, and all-,*B*, including notable representatives like beta barrels, membrane proteins, and enzymes. Those are detailed in Table SI 1.

Before processing and performing the analysis, we preprocessed those 3D structures using pdbfixer^51^ to address nonstandard residues and missing atoms. For our analysis, we excluded irrelevant heteroatoms from each file.

### Simplicial Filtration

Associating simplicial complexes (and indeed a topological space) to a protein structure is a fundamental step. There are numerous ways to achieve this but herein we employ simplicial filtration method to construct the so called Vietoris-Rips complex,^52, 53^ (or equivalently clique complex). For completeness, we provide the basic concepts and specifically how we implement it. A simplicial complex (SC), *K*, is a subset of the power set 2^*V*^ of the set of nodes (vertices) *V* (in the sense of graph theory), with elements of *K* denoted faces (*f* ) and *K* endowed with the hereditary property: given an element *fϵK* and another element *f*′ *⊆f*, then *f*′ *K*. Thus, given an n-dimensional *f* (i.e. a subset of *V* composed of *n* +1 nodes) then its power set 2^*f*^ is called an n-simplex (i.e. n-dimensional polytope being the convex closure of its *n* +1 vertices). If *d* is the maximum dimension of the faces in *K*, then *K* is *d*-dimensional and it is called a simplicial *d*-complex. The underlying graph *K*^1^ of *K* is the graph induced by the 1-simplices in *K* (i.e. the graph whose node set is *V* and whose edge set is the set of all the 1-faces in *K*). Consequently, an n-simplex induces a (n+1)-clique made of 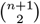edges (i.e. 1-faces) in *K*^*1*^ . In other words, one can associate a simplicial complex with a network by considering a *clique complex*: each clique (complete subgraph) of *n* +1 vertices of the network is seen as an *n*-simplex.

#### Vietoris-Rips complex

given a point cloud or set of vertex *V*⊆ ℝ^*m*^ (e.g. the set of atoms or residues of a protein) then ∀*v* ∈*V* consider the open ball *B*_*ε*_(*v*) ={ *x ∈*ℝ^*m*^ |‖x− ‖ *<ε* }of center *v* and radius *ε>* 0 (i.e. cutoff parameter). Then, a Vietoris-Rips complex *K* has *n*-simplices (i.e. collection of n+1 nodes), {*v*_*i*_*}*⊆*V*, whose balls only require pairwise intersection to be nonempty, that is,

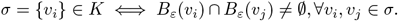

This is equivalently a *clique complex* of a graph with *V* as set of vertices and an edge (*v*_*i*_, *v*_*j*_) if and only if ‖ *v*_*j*_. − *v*_*i*_ ‖ *<* 2ε.

A *finite filtration* that constructs a Vietoris-Rips complex generates a nested sequence of subcomplexes *K*^*i*^

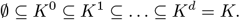

That is, for each 0 ≤ *i*≤*d*, we construct *K*^*i*^ = *K*^*i−*1^ ∪ σ*-*_*i*_, where σ*-*_*i*_ is the set of all *i*-simplices. This leads to *K* ={σ*-*_0_, σ*-*_1_, σ*-*_2_,…, σ*-*_*d*_*}*, which is parameterised by the cutoff parameter.

For our specific implementation, after parsing the protein file (from a database), an initial set of nodes (e.g. atoms or residues) and edges (1-simplex) are determined, where the edges have an assigned weight based on a distance measure between nodes. The minimal and maximum permissible distance between nodes are *a priori* defined. The input to the algorithm is the set of edges and iteratively, higher-order simplices (or cliques) are constructed by adding edges in increasing order of their distance (or weight) with the condition that the *diameter* of the generated simplex (clique) does not exceed a certain bound, where the *diameter* of a clique is the maximum (or minimum) weight of an edge in a clique. Specifically, we bound the computations to *d* = 13 and range the filtration cutoffs between 2.5 to 7.5 Å. Moreover, the edges representing a distance of less than 2 Å were excluded from our graph. To list all the cliques we use the `NetworkX` software package.

### Euler Characteristics and Local Euler Curvature

The *Euler characteristic* (EC), denoted by *x*, is defined as the alternating sum of the number of simplexes of increasing dimensions in the SC. In other words, *x* = *v*_0_*-v*_1_+*v*_2_ *v*_3_+.. .+*v*_*d*_, where *v*_*k*_ is the number of simplices of dimension *k* in the complex and *d* is the dimension of the highest-dimensional simplex in the complex. Noteworthy, the EC is a topological invariant, which means it remains unchanged regardless of how the SC is deformed, as long as the underlying topological structure is preserved. Interestingly, EC can be decomposed (partitioned)^40–42^, which gives a formula for EC via a partition of χ into local components κ_*k*_ (denoted local curvature) for all *k* in the vertex set *V* . Indeed, any scalar quantity which adds up to the Euler characteristic is called curvature. Specifically,

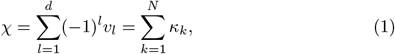

where 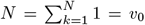 is the cardinality of *V*, that is, the number of nodes of the SC, *v*_*l*_ is the number of simplices of dimension *l* that contain the node *k*, and the *Local Euler Curvature (LEC)* is defined as follows

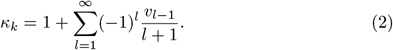

Computing the LEC as per Eq. (1) involves three primary steps: constructing the network, creating the simplicial complex with a specified cutoff, and finally, evaluating curvature κ. We have included an example of simplical complexes in Fig. SI 1, where we compute the respective Euler Characteristics.

Herein, we focus on computing local curvatures, κ_*k*_, of residues. Specifically, given a residue *R* (with a given set of atoms) its local curvature is summed as follows

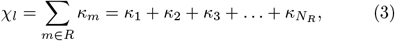

where κ_*m*_ is the local curvature of node *m* of the protein residue *R* network composed on *N*_*R*_ nodes.

After preprocessing, the heavy restricted backbone network, comprising PDB atoms CA, C, N of each protein residue, was created for each 3D structure. Hereafter, the network used to compute the simplical simplex is formed by its restricted backbone and the LEC of a residue is given by PDB atoms CA, C and N of the respective protein, and the LEC of a residue is given by

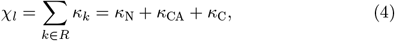

This choice of network makes our results easily interpretable and more accurate. Following a filtration process, then Eq. (1) is used to compute *κ*for each network node. Note that the inter-residue connections were prioritized, while intra-residue connections were disregarded. The LEC was subsequently computed for each residue by summing the *κ*of its corresponding heavy backbone atoms, as described in Eq. (4). The training consensus set average and standard deviations are depicted in Fig. 2 for *α*-Helix,*B*-Strand, and Coils/Turns. Additionally, we show in Figs. SI 3 and SI 4 the full panel comparing DSSP, STRIDE and consensus, as well as, the classification of LEC profiles.

For each residue and respective LEC, we extracted 51 features from the simplicial complexes with cutoffs ranging from 2.5 to 7.5 Å in steps of 0.1 Å. This list of LEC at different cutoffs forms our core feature space for the machine learning algorithm. All data processing utilized in this study are publicly available; see *Data availability* and *Code availability*.

### Unsupervised Clustering

We examined 3 subcategories of the CATH data set: all, consensus only and helices and,*B*-structures (which we will refer to as the *H,B* category). For each category (i.e. those on which the DSSP and STRIDE classifications agree), we assigned to each path a vector in 51-dimensional space, corresponding to the LEC calculated for the protein at cutoffs from 2.5 Å to 7.5 in steps of 0.1 Å.

To prepare the data for clustering, we performed dimensionality reduction via principle component analysis (PCA). PCA is a common procedure used for basic dimensionality reduction and denoising, which projects the data onto linear subspaces on which the data has the largest variance, thus preserving the significant features which theoretically getting rid of extra dimensionality occurring due to noise. We performed PCA using the inbuilt Scikit-learn class^54^, which indicated that just above 75% of the variance in the data was contained in the first 5 PCs in the case of the consensus category and full CATH data set. The *H,B* category showed even higher correlations, with 75% of the variance explained by the first 2 PCs.

We chose to project the data to the first 5 PCs as the gains in explained variance ratio rapidly decreased after this point. For clustering, we fit a series of Gaussian Mixture Models (GMMs). GMMs represent a dataset as a mixture of multiple Gaussian distributions with (possibly) different parameters, allowing the model to capture more complex patterns than, for example, a *k*-means based method. We fit GMMs with *n* clusters, *n* ∈{2, 3, 4, 5, 6, 7, 8, 9, 10} directly to the 5-dimensional projection effected by PCA.

We tested the robustness of the clustering using a variety of measures. The Jensen-Shannon metric^55^ quantifies the dissimilarity or divergence between two probability distributions. As GMMs are essentially probability distributions, the Jensen-Shannon metric can be used to measure the similarity of GMMs. By fitting differing GMMs on subsets of the data and measuring the similarity of the models, we can test the stability of the clustering.

Similarly, we can measure the homogeneity and completeness of GMMs trained on different subsets of the data. Homogeneity measures the degree to which each cluster contains only data points that are members of a single class. A high homogeneity score indicates that the clusters are consistent with the data categories, which is desirable in clustering. Completeness quantifies the degree to which all data points that are members of a particular class are assigned to the same cluster. In essence, it evaluates whether all data points of a given class are correctly clustered together. A high completeness score indicates that the clustering algorithm successfully captures all data points from the same category within a single cluster. By comparing the homogeneity and completeness of clusters of one GMM to the classes assigned by another, we can test how much much the cluster predictions would change when starting with a different sample class. See Fig. SI 9 for results.

The consensus categories and full CATH categories gave broadly similar results. The metrics and clustering of the *H,B* set was markedly differentiated; see Fig. SI 10.

### Random Forest Classifier

Our ensemble of random trees was created by using Scikitlearn^54^ (sklearn) with 100 trees. They were fitted using a balanced class-weight and an out-of-bag strategy was used to assess the accuracy of the predictors. The tree minimum amount of samples to split a leaf was exhaustively explored using a nested resampling strategy. The cross-validation used an inner and outer resampling of 5 folds, both resamplings were stratified and shuffled. For each fit, the accuracy, balanced accuracy, f1-weighted and f1-macro scores were collected and they are shown in Fig. SI 13a. Subsequently, the classification model was trained using the CATH data set of consensus classification, using a minimum of 100 samples to split each leaf. The final validation scores, from the test data set, is also shown in Fig. SI 13a, where we selected our hyperparameter at approximately the top of the local maximum of f1-macro score.

Our training set includes only the residues where the DSSP and STRIDE classification agree, namely, the consensus classification. All other residues were left to check the performance of our the model. The final scores are described into Table 1, and Table SI 2 shows the confusion matrix resulting from the consensus classification. Detailed additional scores for the test set are described in Table SI 1. More technical information is available in the Supporting Information, Section 2.

## Data availability

Data of LEC from CATH is public available at DOI:10.5281/zenodo.8382584^49^. Data of LEC from test set at Table SI 1 and Random Florest Classifier and respective Python3 Pickle model are public available at DOI:10.5281/zenodo.8385504. Python Jupyter-notebooks from Unsupervised Clustering are available at DOI:10.5281/zenodo.8385504. The data is distributed under CC BY-NC-ND 4.0 (https://creativecommons.org/licenses/by-nc-nd/4.0/) license.

## Code availability

Source codes related to LEC processing and Results in this paper are public available at DOI:10.5281/zenodo.8382584^49^ and DOI:10.5281/zenodo.8385504. The source codes are distributed under CC BY-NC-ND 4.0 (https://creativecommons.org/licenses/by-nc-nd/4.0/) license. Alternatively, the codes are available for download via the following github repository: https://github.com/MCEN-BCAM/LEC_Protein-SecondaryStructures.

## Acknowledgements

This research is supported by the Basque Government through the BERC 2022-2025 program and by the Ministry of Science and Innovation: BCAM Severo Ochoa accreditation CEX2021-001142-S/MICIN/AEI/10.13039/501100011033. We also acknowledge financial support received from the IKUR Strategy under the collaboration agreement between Ikerbasque Foundation and BCAM on behalf of the Department of Education of the Basque Government. Moreover, we thank the Basque Modelling Task Force (BMTF) grant from the Basque Government. All authors thank Prof. Oliver Knill for useful discussions.

## Author contributions statement

R.A.M. conceived the paper, the LEC formulation and Random Forest Classifier. R.B. contributed to the unsupervised clustering and associated results. S.R. and I.U.-B. supervised the project, acquired the funding. S.R conceptualised some of the ideas. M.D. contributed to the funding. All authors contributed to the writing of the manuscript.

## Competing interests

The authors declare no competing interests.

## Additional information

To include, in this order: **Accession codes** (where applicable); **Competing interests** (mandatory statement).

The corresponding author is responsible for submitting a competing interests statement on behalf of all authors of the paper. This statement must be included in the submitted article file.

## Notes

https://zenodo.org/records/8382584

https://github.com/MCEN-BCAM/LEC_ProteinSecondaryStructures

### Competing Interest Statement

The authors have declared no competing interest.

