## Supporting Information document for "Discovering Secondary Protein Structures via Local Euler Curvature"

[1]Basque Center for Applied Mathematics, 48009 Bilbao, Bizkaia (Basque-Country, Spain)
[2]Dutch Institute for Emergent Phenomena (DIEP), Institute for Advanced Study (IAS) and University of Amsterdam, 1012 GC, Amsterdam, The Netherlands
[3]Department of Medicine, Division of Geriatrics, Faculty of Medicine and Health Sciences, Universit de Sherbrooke, Sherbrooke, QC, Canada
[4]Research Center on Aging, Centre Intégré Universitaire de Santé et Services Sociaux de l'Estrie-Centre Hospitalier Universitaire de Sherbrooke, Sherbrooke, QC, Canada
[5]Inria Centre at Université Côte d'Azur, 06902 Sophia Antipolis cedex, France
[6]Université Côte d'Azur, 06103 Nice cedex 2, France
[7]Instituto Biofisika (UPV/EHU, CSIC), University of the Basque Country, E-48940, Leioa, Spain
[8]Ikerbasque, the Basque Science Foundation, Bilbao, Spain
[*]
[†]

**This Supporting Information expands the information about our unsupervised clustering and the Random Forest classifier model used in the main text. It also list additional Figures, Tables and References.**

Keywords: Computational Topology, Euler Characteristics, Local Euler Curvature, Protein Structure, Random Forest Classifier, Unsupervised Clustering

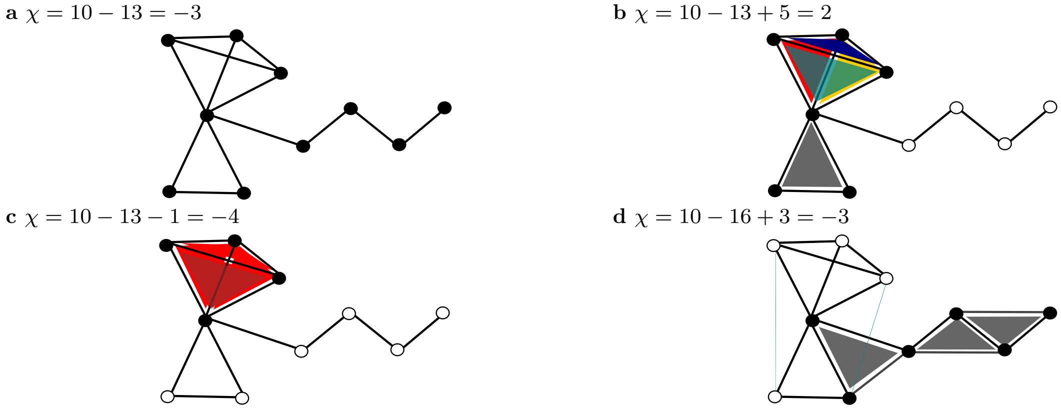


**Fig. SI 1. Examples of simplicial complexes.** **(a)** A SC composed of ten 0-simplex (nodes) and thirteen 1-simplex (edges). **(b)** Five 2-simplex (triangles). **(c)** One 3-simplex (tetrahedron) from **a**. **(d)** Adding more edges we get three new 2-simplexes nonexistent in **b,c**. The Euler Characteristics are explicitly given for each SC.

#### Local Euler Curvature

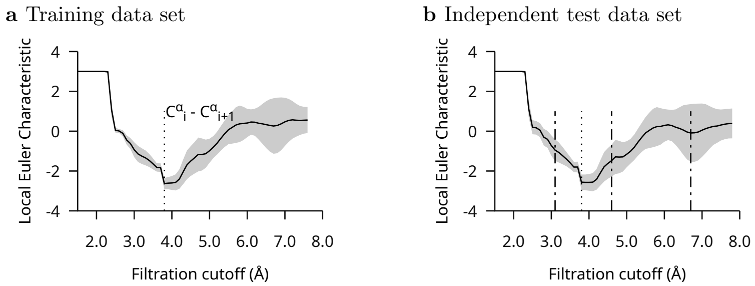


**Fig. SI 2.** Caption

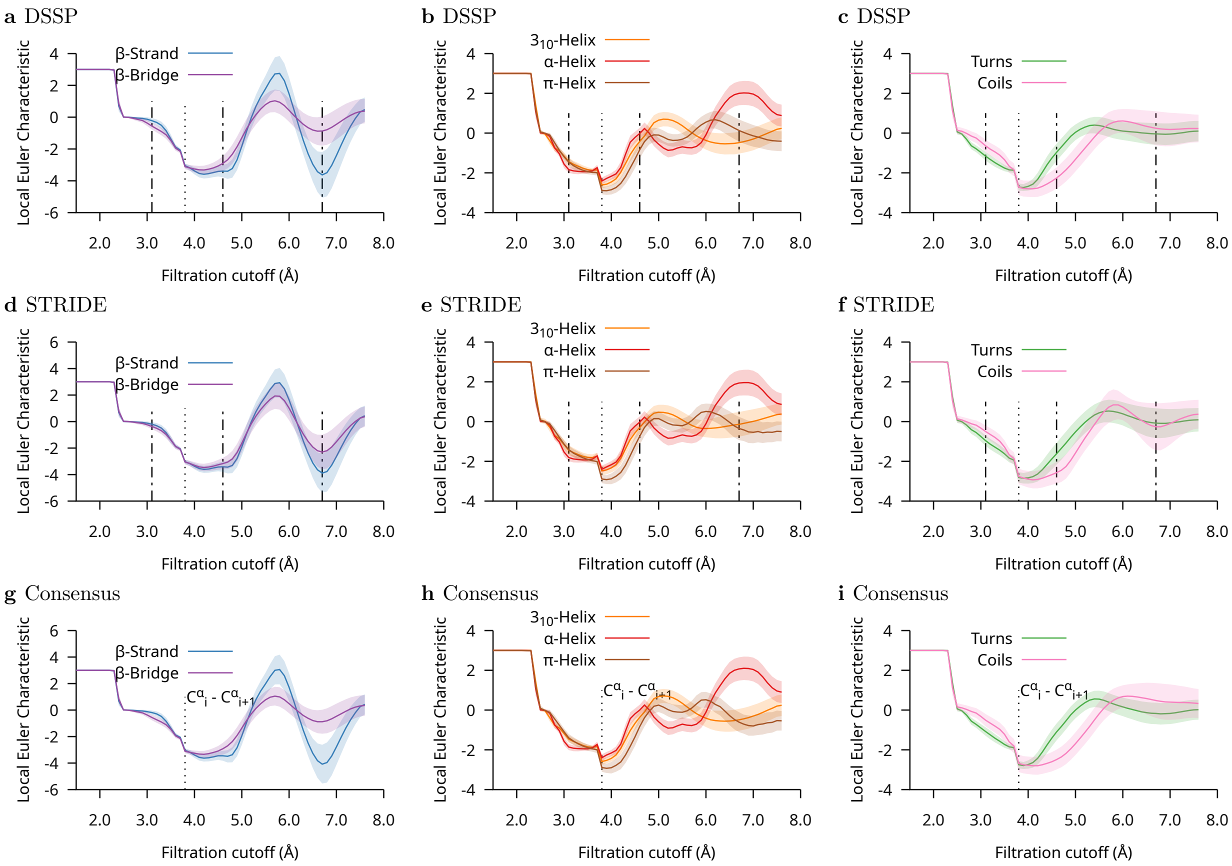


**Fig. SI 3.** CATH data set analysis. Average of Local Euler Curvature of residues considering DSSP, STRIDE and the consensus (STRIDE = DSSP) assignment. The shaded area represents the standard deviation.

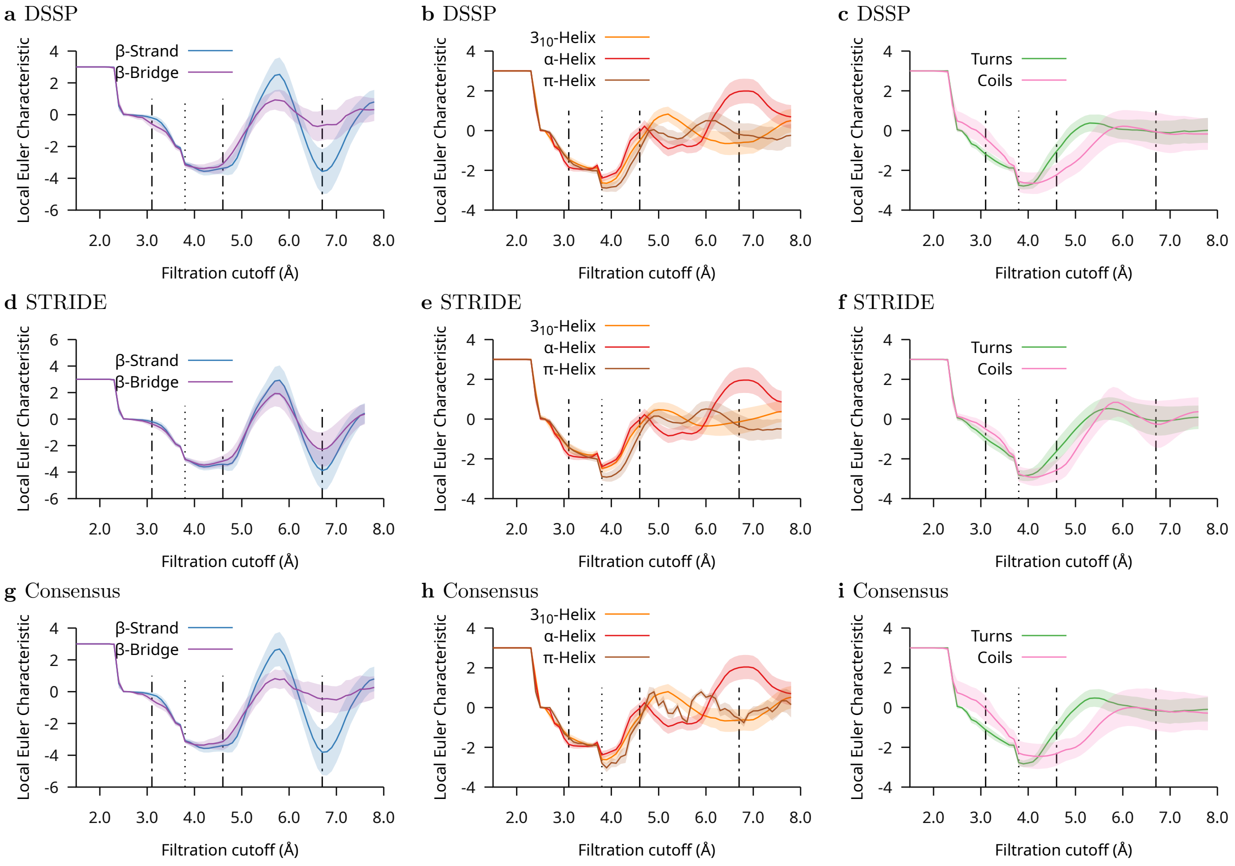


**Fig. SI 4.** Independent test data set analysis. Average of Local Euler Curvature of residues considering DSSP, STRIDE and the consensus (STRIDE = DSSP) secondary structure assignments. The shaded area represents the standard deviation.

### Unsupervised Clustering of LEC profiles

#### Principal Component Analysis (PCA)

Principal Component Analysis (PCA) plays a pivotal role in our research as a powerful dimensionality reduction technique. In the study of protein structures, PCA assists in simplifying the complex multidimensional space represented by the LEC profiles. These profiles, which capture essential information about the spatial shape of proteins, often exist in high-dimensional spaces due to numerous data points. PCA allows us to distill this intricate information into a smaller set of orthogonal components known as Principal Components (PCs). The first few PCs typically account for the majority of the variance within the dataset, making them valuable for data compression and visualization. In our analysis, PCA serves as an essential step in reducing the dimensionality of LEC profiles while preserving their structural essence, enabling subsequent clustering and classification processes.

PCA's role extends beyond dimensionality reduction; it also facilitates pattern recognition and data exploration in our research. By projecting data points onto the subspace defined by the top PCs, we can identify key structural features and relationships among protein structures. This helps in uncovering hidden patterns and differences that might not be immediately evident in the original high-dimensional data. Moreover, PCA aids in identifying the principal directions along which structural variations occur, potentially revealing insights into the evolutionary or functional aspects of protein structures. In essence, PCA serves as a foundational tool that empowers us to navigate the intricate landscape of protein structural data, simplifying it for further analysis while preserving its critical information content.

#### Gaussian Mixture Models (GMMs)

Gaussian Mixture Models (GMMs) constitute a crucial component of our research methodology for deciphering the structural complexities of protein ensembles. In the context of our study, GMMs serve as a powerful probabilistic model to capture the underlying distribution of protein conformations within identified clusters. When dealing with protein structural data, GMMs enable us to express the dataset as a combination of multiple Gaussian distributions, each representing a distinct structural archetype or cluster. This modeling approach is particularly apt for handling the inherent heterogeneity in protein structures, where a single Gaussian distribution may not adequately represent the diverse conformational space.

By employing GMMs, we can identify the optimal number of clusters, each characterized by its mean and covariance matrix, which corresponds to a specific protein structural pattern. This statistical approach not only aids in clustering similar protein structures but also provides probabilistic membership estimates, indicating the likelihood of a particular protein conformation belonging to each cluster. This probabilistic assignment is particularly valuable in cases where protein structures exhibit overlap between clusters or gradual transitions. Overall, GMMs facilitate the unsupervised clustering of protein structures, contributing significantly to our ability to uncover hidden structural relationships, identify unique conformational patterns, and gain deeper insights into the diverse world of protein structural biology.

#### Measuring the robustness of clustering

The Jensen-Shannon metric was calculated by training a 2 GMMs on two sets of 50% of the proteins selected randomly from each category data set (consensus, full and $H\beta$).

To measure the consistency among clusterings we trained a GMM on 20% of the PCA reduced LEC profiles of proteins from the subset of the CATH data set associated to the relevant category with $n$ clusters for $n \in \{2, ..., 9\}$. In order to estimate the robustness of the resulting classification we then selected 5 more randomly selected subsets representing 20% of the data sets and retrained 50 GMMs (5 each with number of clusters being 2 through 9). Finally, we evaluated the resulting predicted labels for homogeneity and completeness. The average scores over $k$ runs are given in the table. 4 and 6 clusters display better homogeneity and completeness scores for both the consensus and total data sets.

#### UMAP Algorithm

Uniform Manifold Approximation and Projection (UMAP)[13] was used to visualise the relationships between the clusters of LEC profiles and the LEC profiles of proteins defined by the data sets. UMAP operates on the principle of manifold learning, aiming to faithfully represent high-dimensional data in lower-dimensional spaces while preserving the local and global relationships between data points, making it particularly useful at visualising the interrelations of clusters of LEC profiles.

#### Additional Figures and Tables

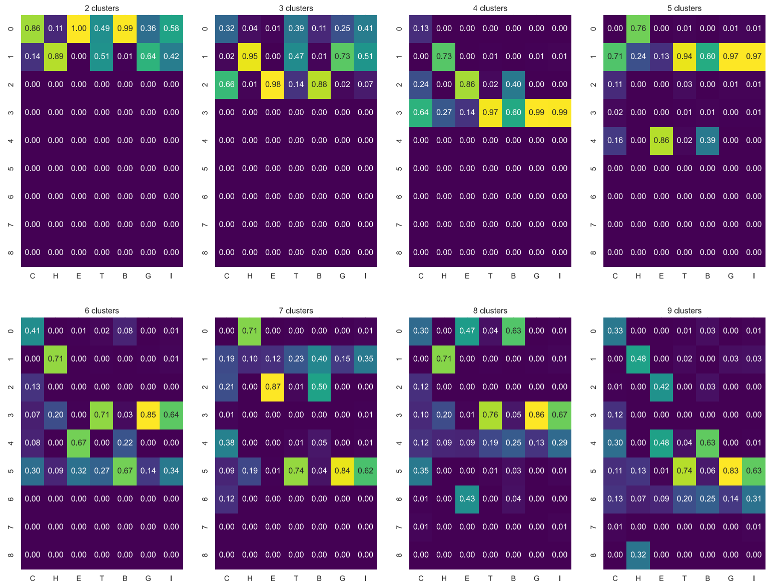


**Fig. SI 5.** Interaction of clustering and consensus labels on the CATH data set.

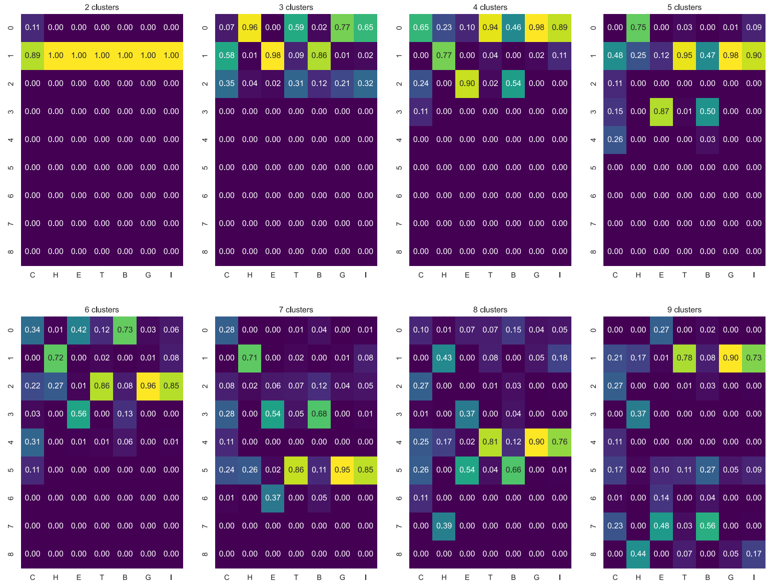


**Fig. SI 6.** Interaction of clustering and DSSP labels on the CATH data set.

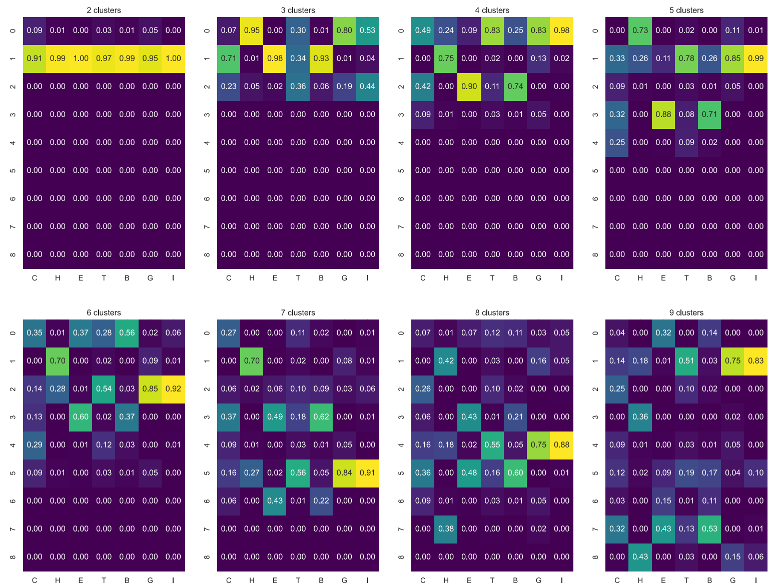


**Fig. SI 7.** Interaction of clustering and STRIDE labels on the CATH data set.

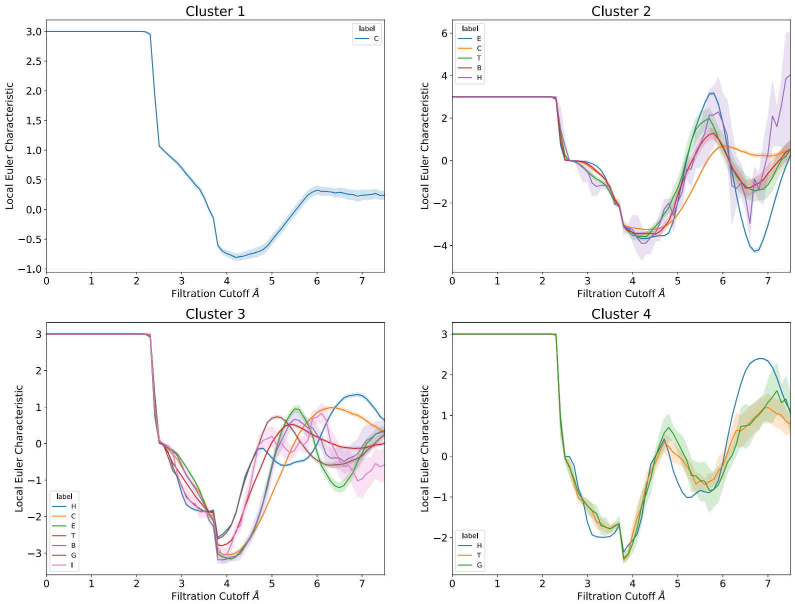


**Fig. SI 8.** Curves with different DSSP and Stride labels can demonstrate remarkably similar LEC profiles.

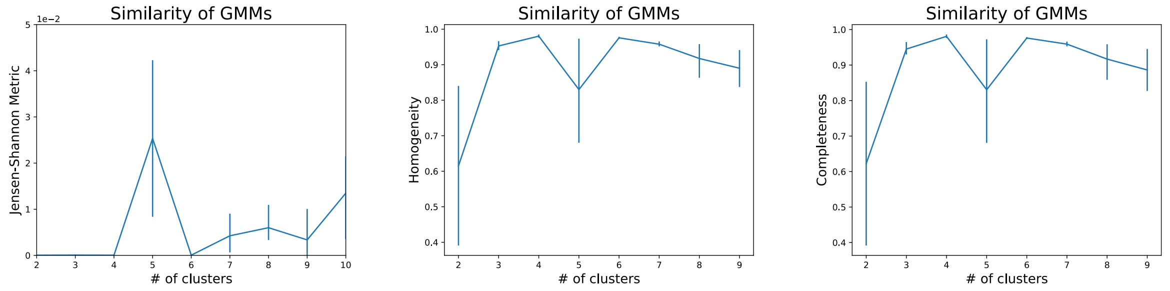


**a** Jensen-Shannon metric **b** Homogeneity of GMM predicted labels. **c** Completeness of GMM predicted labels.

**Fig. SI 9.** Measuring the robustness of clustering, see Sec. Measuring the robustness of clustering for details.

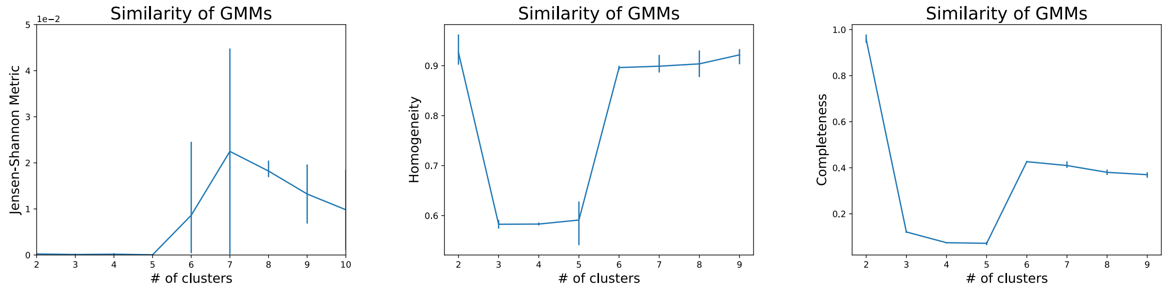


**a** Jensen-Shannon metric **b** Homogeneity of GMM predicted labels. **c** Completeness of GMM predicted labels.

**Fig. SI 10.** Measuring the robustness of clustering, see Sec. Measuring the robustness of clustering for details.

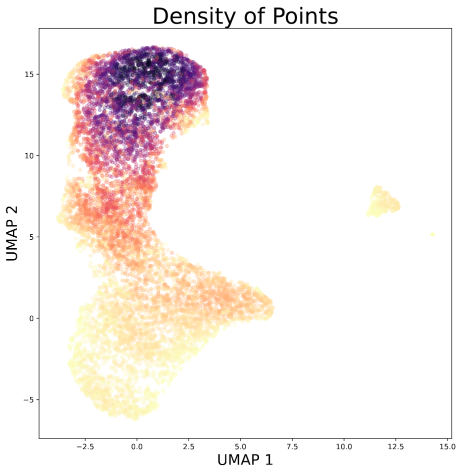


**a** UMAP visualistion with density.

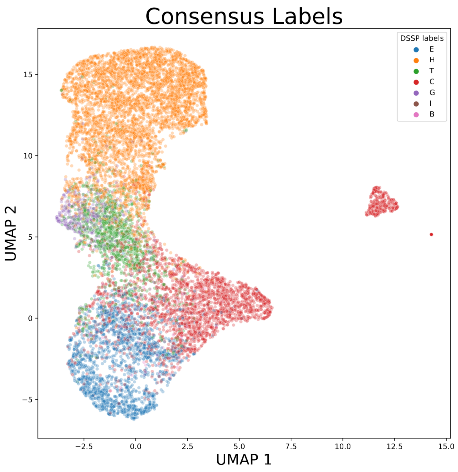


**b** UMAP visualistion with consensus labels.

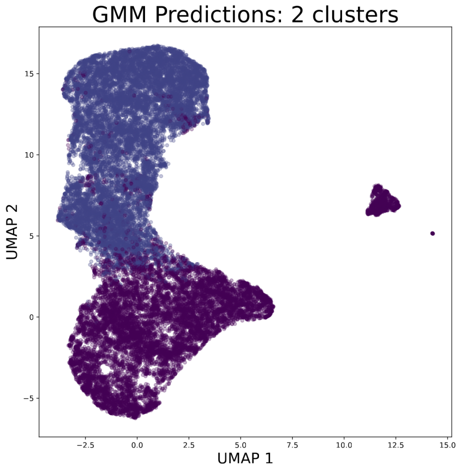


**c** UMAP visualisation with predicted GMM labels.

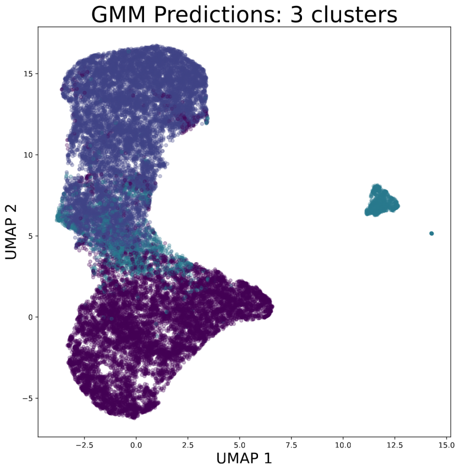


**d** UMAP visualisation with predicted GMM labels.

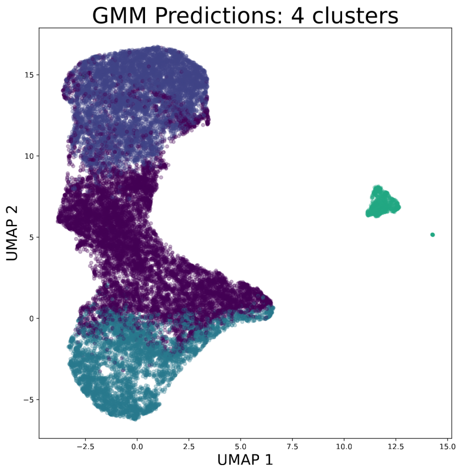


**e** UMAP visualisation with predicted GMM labels.

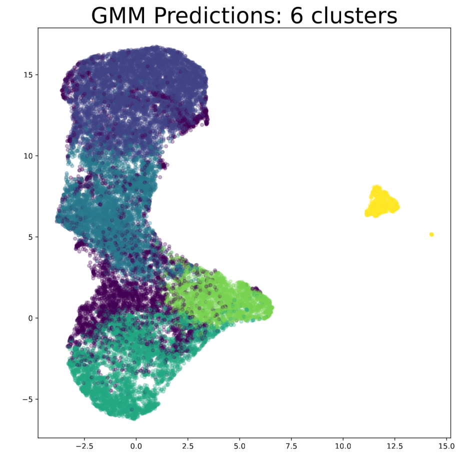


**f** UMAP visualisation with predicted GMM labels.

**Fig. SI 11.** Visualisation of the vectors associated to the LEC profiles of the CATH data set

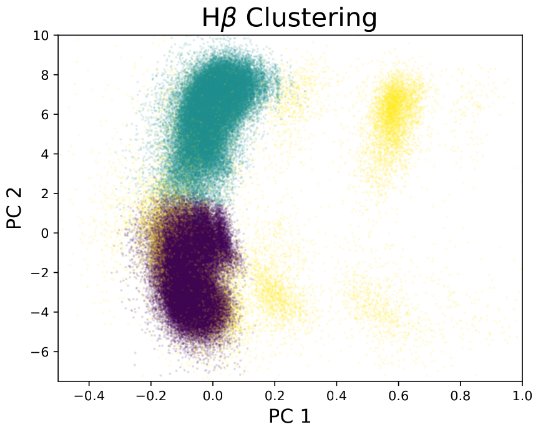


**a** PC components of LEC profiles given by proteins from the $H\beta$ set identified by GMMs trained on the $H\beta$ set

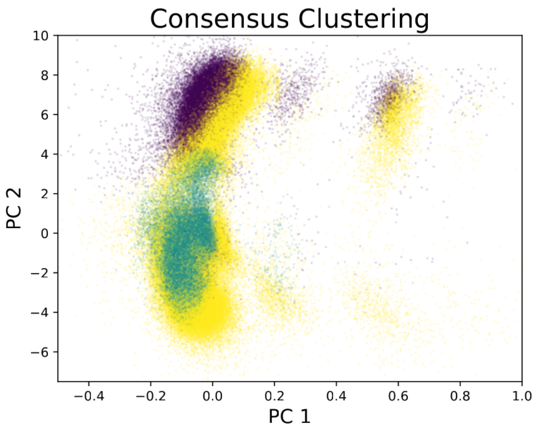


**b** PC components of LEC profiles given by proteins from the $H\beta$ set identified by GMMs trained on the consensus set

**Fig. SI 12.** Although clustering on the consensus set can broadly capture some behaviour in the $H\beta$ sets, $H\beta$ demonstrates its own remarkable clustering patterns unseeable from the full consensus set.

#### Random Forest

We are poised to demonstrate the remarkable potential of the Euler Characteristic (EC) as a versatile and effective tool for elucidating the intricate structures of proteins. To substantiate its efficacy, we employ a straightforward yet powerful machine learning (ML) algorithm, the Random Forest[2], for classifying protein structures based on DSSP classification.

The Random Forest algorithm, a member of the ensemble learning family, capitalizes on the collective strength of multiple decision trees to enhance predictive performance while mitigating overfitting. During training, it crafts a multitude of decision trees, with each tree honed on a random subset of the data and using a random subset of features. These individual trees independently make predictions, and the final decision materializes through majority voting. By amalgamating predictions from diverse and decorrelated trees, the Random Forest mitigates variance, enhances accuracy, and bestows resilience against data noise and outliers[5].

The Random Forest's ability to adeptly navigate high-dimensional data while retaining interpretability for feature importance analysis has rendered it indispensable in a myriad of domains, encompassing bioinformatics[18], noisy datasets[20], applied mathematics[1], health sciences[17,21,29], and beyond[10,22].

Our intuitive approach involves adopting the Local Euler Curvature (LEC) filtration as a feature space. Subsequently, our cutoff distances span the range from 2.0 to 7.5 Å in 0.1 Å increments. In our modeling, each residue serves as a sample, characterized by 55 features that correspond to the filtration of LEC, as defined in Eq. 4. This model bears the moniker **rbLEC**, signifying 'Restricted Backbone LEC'.

Decision Trees[2], a widely embraced machine learning algorithm for classification and regression tasks, operate through recursive partitioning of data into subsets. This process is primarily driven by informative sample features, constructing a tree-like structure. Internal nodes represent features, while leaf nodes correspond to anticipated class predictions or regression values. The decision-making journey commences at the root node and progresses down the tree, evaluating feature conditions until reaching a leaf node for prediction. The core objective of this partitioning process is to optimize information gain or reduce impurity within subsets, ensuring that the resultant tree effectively captures data patterns and relationships.

To establish a robust training set, we employ the same CATH database of non-redundant protein structures as previously. This selection deliberately includes consensus residues, further enhancing the separability of the feature space (see Fig. SI 3). For comprehensive insights into the model's hyperparameters, refer to the Methods section.

Our trained model also facilitates an assessment of the relative importance of each feature within our filtration protocol. Building upon our prior analysis showcased in Fig. 2, which identified specific cutoff distances where LEC averages and standard deviations are presumed to influence secondary structure classification, we employ a decision tree-based rationale rooted in the general shapes of these LEC paths to distinguish among diverse structures. Consequently, we validate this reasoning by scrutinizing the computed relative importance of each feature. Fig. 3b unveils three prominent peaks at 3.1, 4.6, and 6.7 Å. In harmony with our observations across Fig. 1d-1f, these cutoff distances offer expressive representations of $\chi_k$ pertaining to 2GB1's secondary structures. Moreover, in Fig. 2c, we witness the 3.1 Å range effectively segregating all seven classes, while at 4.6 Å, a discernible separation emerges between $\alpha$ and $\beta$ structures. The final significant feature, the 6.7 Å cutoff, facilitates differentiation between $\alpha$-Helix, and $\pi$-Helix/$3_{10}$-Helix, (Fig. 2e) while also distinguishing $\beta$-Strand, from $\beta$-Bridge, as seen in Fig. 2d.

Hence, our earlier training analysis seamlessly aligns with the observed relative feature importance post-training of the random forest classifier. This congruence underscores the representation of training ensemble averages in Fig. 2 and validates the authenticity of the LEC paths' representation of each secondary structure class.

Furthermore, we subject our model to rigorous testing through an independent assessment test set (Table SI 1). The post-training results manifest as impressive scores of 0.87, 0.88, 0.65, and 0.70 for accuracy, f1-weighted, balanced accuracy, and f1-macro, respectively. Fig. SI 13a aptly illustrates the remarkable concordance between these scores and our nested cross-validation protocol.

**Table SI 1.** Scores relative of protein Data sets used for final assessment. We show the number of residues and detailed scores as obtained after our trained random forest ensemble.

| Residues | | f1-score | | | | Accuracy | |
|---|---|---|---|---|---|---|---|
| PDB | #res | $\beta$-Strand | $\alpha$-Helix | Turn | Coil | Score | $\text{Top}_{K=2}$ |
| 108L[4] | 164 | 0.75 | 0.97 | 0.63 | 0.86 | 0.91 | 0.95 |
| 1DEH[8] | 748 | 0.92 | 0.91 | 0.68 | 0.80 | 0.83 | 0.93 |
| 1DUB[9] | 1566 | 0.88 | 0.96 | 0.57 | 0.84 | 0.85 | 0.95 |
| 1EMA[15] | 225 | 0.96 | 0.67 | 0.80 | 0.90 | 0.88 | 0.96 |
| 1K1X[12] | 1318 | 0.93 | 0.95 | 0.73 | 0.86 | 0.88 | 0.94 |
| 1MJ5[14] | 302 | 0.87 | 0.97 | 0.64 | 0.90 | 0.87 | 0.94 |
| 1PFK[23] | 640 | 0.91 | 0.96 | 0.74 | 0.86 | 0.90 | 0.94 |
| 1RBP[7] | 182 | 0.90 | 0.88 | 0.74 | 0.83 | 0.82 | 0.91 |
| 1SYN[26] | 528 | 0.93 | 0.91 | 0.60 | 0.86 | 0.83 | 0.94 |
| 2PIL[11] | 158 | 0.80 | 0.94 | 0.72 | 0.84 | 0.84 | 0.91 |
| 7DF4[28] | 4408 | 0.85 | 0.94 | 0.68 | 0.88 | 0.84 | 0.93 |
| 7G87 | 430 | 0.91 | 0.97 | 0.83 | 0.93 | 0.93 | 0.97 |
| 7YM9 | 536 | 0.91 | 0.90 | 0.79 | 0.89 | 0.86 | 0.93 |
| 8AD3 | 556 | 0.91 | 0.97 | 0.79 | 0.86 | 0.89 | 0.97 |
| 8B0N[25] | 532 | 0.91 | 0.94 | 0.65 | 0.89 | 0.89 | 0.94 |
| 8BS3[16] | 456 | 0.91 | 0.94 | 0.77 | 0.92 | 0.90 | 0.96 |
| 8CPH[3] | 554 | 0.88 | 0.96 | 0.64 | 0.86 | 0.90 | 0.96 |
| 8DYJ | 762 | 0.89 | 0.96 | 0.74 | 0.93 | 0.92 | 0.96 |
| 8FN8[24] | 337 | 0.89 | 0.92 | 0.79 | 0.86 | 0.86 | 0.94 |
| 8GQC[19] | 264 | 0.93 | 0.94 | 0.68 | 0.85 | 0.83 | 0.96 |
| 8IUM[27] | 1397 | 0.88 | 0.97 | 0.78 | 0.85 | 0.87 | 0.96 |
| 8T1O[6] | 1812 | 0.90 | 0.95 | 0.68 | 0.93 | 0.90 | 0.95 |

**Table SI 2. Confusion matrix from consensus assignment of protein secondary structures.** The four major classes, $\beta$-Bridge, $\beta$-Strand, $\alpha$-Helix and Turns Coils, represent roughly 95% of all 13635 residues/samples available. All test data is described in Table SI 1.

| | $\beta$-Bridge | $\beta$-Strand | $3_{10}$-Helix | $\alpha$-Helix | $\pi$-Helix | Turns | Coils |
|---|---|---|---|---|---|---|---|
| $\beta$-Bridge | 22 | 24 | 0 | 0 | 0 | 5 | 27 |
| $\beta$-Strand | 123 | 2558 | 0 | 0 | 0 | 71 | 204 |
| $3_{10}$-Helix | 1 | 0 | 397 | 7 | 0 | 86 | 0 |
| $\alpha$-Helix | 4 | 0 | 236 | 4955 | 3 | 216 | 21 |
| $\pi$-Helix | 0 | 0 | 1 | 0 | 2 | 2 | 0 |
| Turns | 23 | 7 | 114 | 30 | 2 | 957 | 92 |
| Coils | 81 | 223 | 2 | 9 | 0 | 170 | 3008 |

**a**

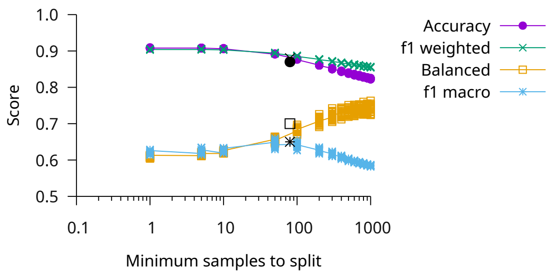


**Fig. SI 13. Random forest classifier. (a)** Validation of our classifier obtained by a nested cross-validate resampling exhaustive exploring the minimum samples to split a leaf of the tree model. The black points are the scores got by the final independent test set.

**21.** K Saikumar and V Rajesh. A novel implementation heart diagnosis system based on random forest machine learning technique. *International Journal of Pharmaceutical Research (09752366)*, 2020.

**22.** Syed Haleem Shah, Yoseline Angel, Rasmus Houborg, Shawkat Ali, and Matthew F McCabe. A random forest machine learning approach for the retrieval of leaf chlorophyll content in wheat. *Remote Sensing*, 11(8):920, 2019.

**23.** Yasuo Shirakihara and Philip R Evans. Crystal structure of the complex of phosphofructokinase from escherichia coli with its reaction products. *Journal of molecular biology*, 204(4):973–994, 1988.

**24.** Ashis Sinha, Jessica Kawakami, Kimberly S Cole, Aliona Ladutska, Mary Y Nguyen, Mary S Zalmai, Brandon L Holder, Victor M Broerman, Russell T Matthews, and Samuel Bouyain. Protein-protein interactions between tenascin-r and rptp$\zeta$/phosphacan are critical to maintain the architecture of perineuronal nets. *Journal of Biological Chemistry*, page 104952, 2023.
